# A range of voltage-clamp protocol designs for rapid capture of hERG kinetics

**DOI:** 10.1101/2024.08.14.607756

**Authors:** Chon Lok Lei, Dominic G. Whittaker, Monique J. Windley, Matthew D. Perry, Adam P. Hill, Gary R. Mirams

## Abstract

We provide details of a series of short voltage-clamp protocols designed for gathering a large amount of information on hERG (K_v_11.1) ion channel gating. The protocols have a limited number of steps and consist only of steps and ramps, making them easy to implement on any patch clamp setup, including automated platforms. The primary objective is to assist with parameterisation, selection and refinement of mathematical models of hERG gating. We detail a series of manual and automated model-driven designs, together with an explanation of their rationale and design criteria. Although the protocols are intended to study hERG1a currents, the approaches could be easily extended and generalised to other ion channel currents.

## 1 Introduction

This report describes a series of voltage-clamp protocol waveforms that were designed to explore the gating of cell lines expressing hERG1a / K_v_11.1 channels, which are the primary subunit of the channels carrying the cardiac rapid delayed rectifier potassium current, I_Kr_ (Sanguinetti et al., 1995; Vandenberg et al., 2012).

The aim is to build on our previous studies that aimed to develop a range of short, information-rich voltage clamp protocols to use in experimental recordings to capture hERG gating behaviour (Beattie et al., 2018; Lei et al., 2019b). Here we extend these to a wide range of protocols to better parameterise, select and test mathematical models of hERG gating (Bett et al., 2011) and in particular to gain a better understanding and quantification of model discrepancy — when models cannot correctly predict what happens in reality (Shuttleworth et al., 2024). As a result, some protocols will focus on classic optimal experimental design in terms of reducing uncertainty / improving identifiability of model parameter estimates (Lei et al., 2023). Whilst others focus on maximising differences between trained models to assist in model selection/discrimination.

All these protocols were designed during the Isaac Newton Institute’s Fickle Heart programme in May–June 2019 (Mirams et al., 2020). The protocols are all designed to be run on an automated patch platform, namely the Nanion SyncroPatch384PE (Obergrussberger et al., 2016), which at the time had a restriction of only allowing up to 64 commands (steps or ramps) to define a single voltage-clamp protocol.

## 2 Models used in protocol design process

Our designs are model-driven akin to Lei et al. (2023), where mathematical models are used as part of automatic optimal design; even where our designs are manual they were done by visually examining the results of forward simulations.

The model structures that we used here are Beattie et al. (2018) and Wang et al. (1997) (also used in Fink et al. (2008)), with their Markov diagrams shown in Figure 1. The first model (Beattie et al., 2018) is a Hodgkin-Huxley style model with two independent gates, which can be represented as a symmetric 4-state Markov model (Rudy and Silva, 2006, Fig. 4B). The second model Wang et al. (1997) is a 5-state Markov model with 3 closed states, an open state, and an inactivated state connected sequentially.

**Figure 1.**
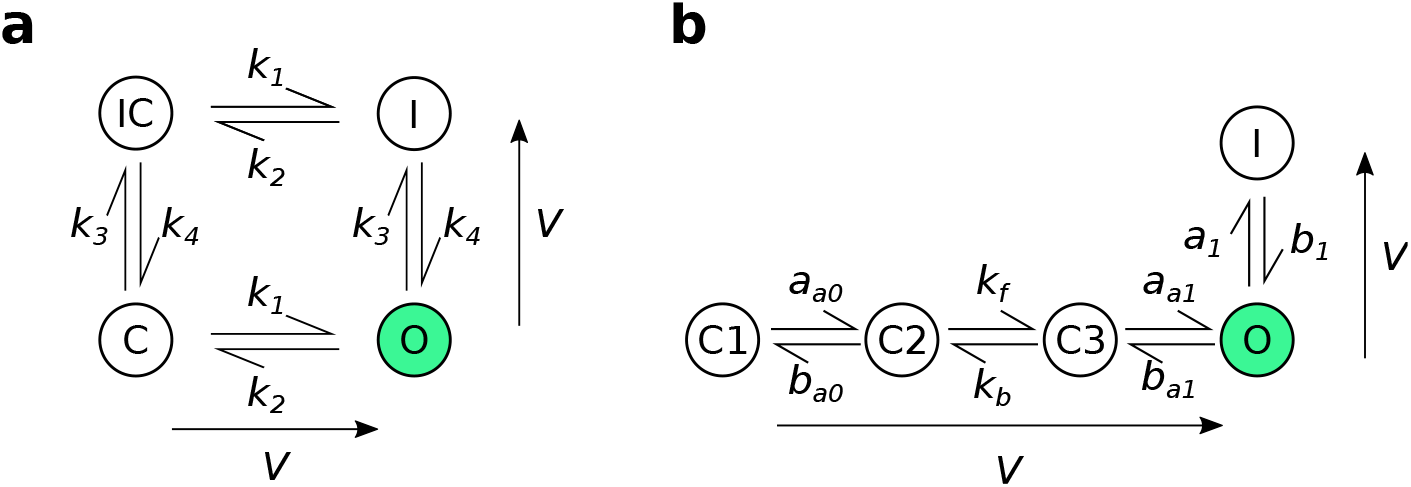
The model structures used for experimental design. (**a**): the four-state Beattie et al. (2018) model. (**b**): the five-state Wang et al. (1997) model. The arrows adjacent to each model structure indicate the direction in which rates increase as the voltage increases. Reproduced from Shuttleworth et al. (2024) under a CC-BY licence.

**Figure 2.**
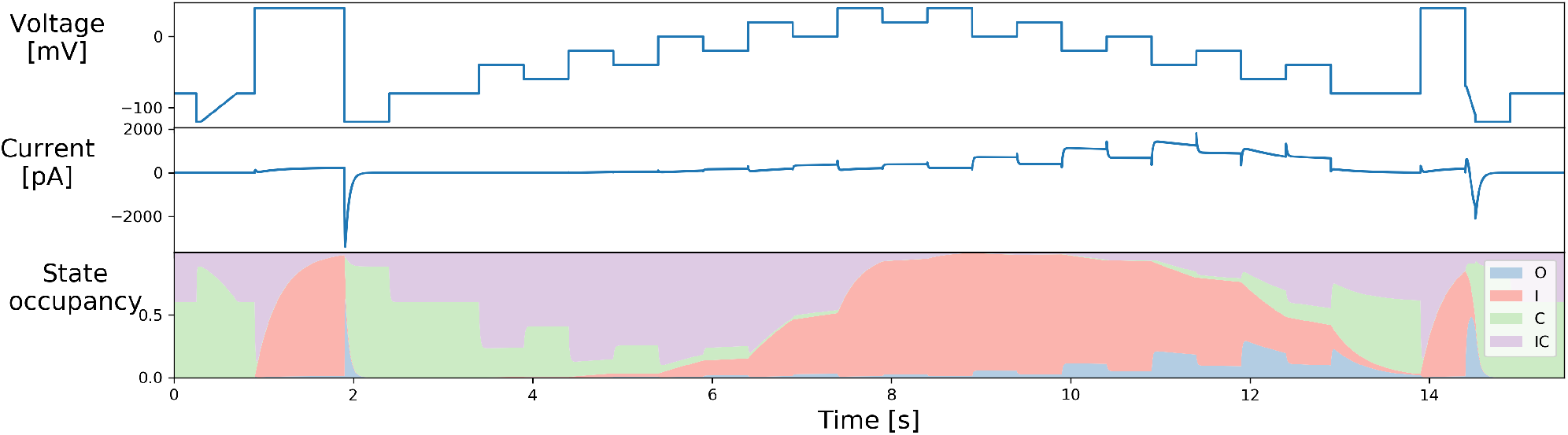
The staircase protocol and its simulation, with state occupancy shown for the Beattie et al. (2018) model of Fig. 1a. Reproduced from (Lei et al., 2019b) under a CC-BY licence.

**Figure 3.**
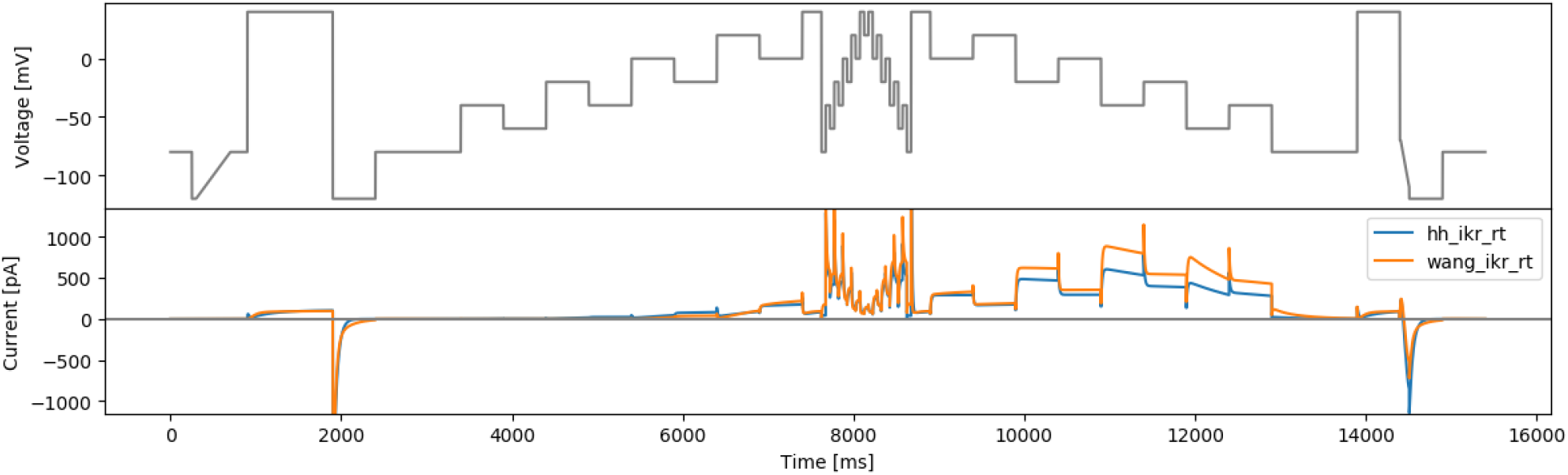
The staircase-in-staircase (sis) protocol and simulated currents from both models. (`hh_ikr_rt` is the Beattie et al. (2018) model of I_Kr_ and `wang_ikr_rt` is the Wang et al. (1997) model of I_Kr_, both parameterised to room temperature data).

**Figure 4.**
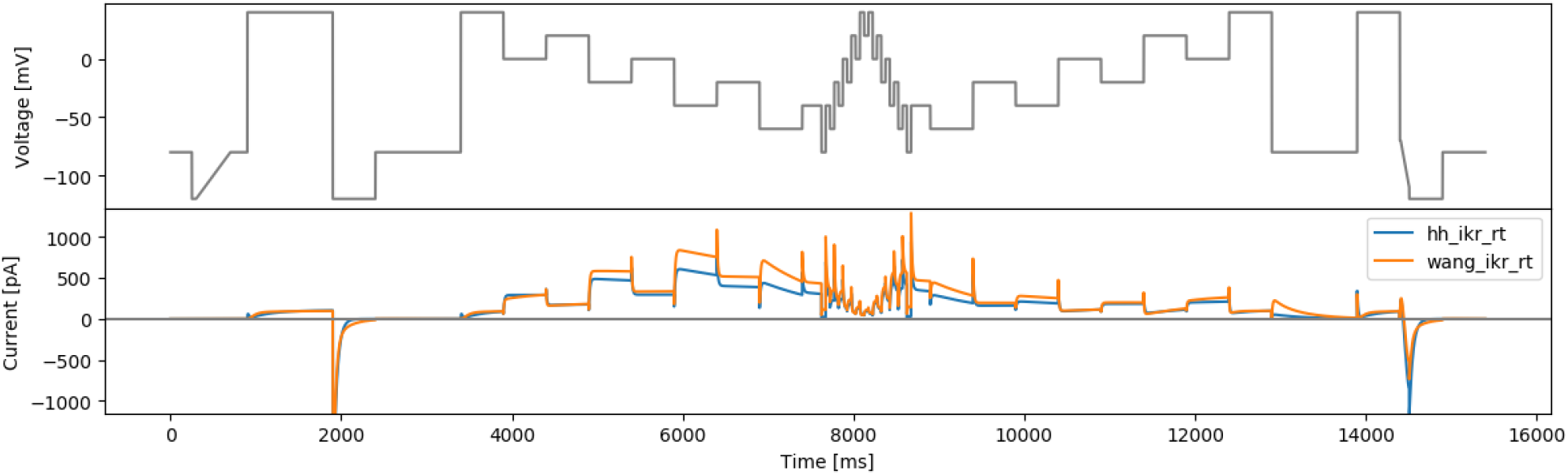
The inverted staircase-in-staircase (sisi) protocol and simulated currents from both models

### 2.1 Beattie model

In matrix/vector form, the Beattie et al. (2018) model can be written as,

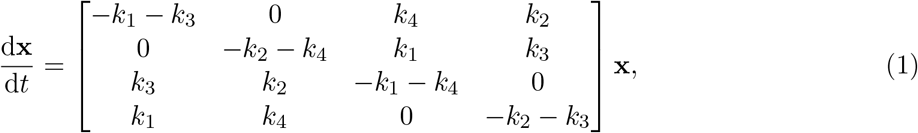

where

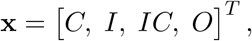

and

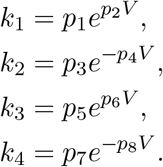

This model is equivalent to a two gate Hodgkin-Huxley style gating model with open probability given by an “activation” *a* gate representing the ‘right’ transitions in Fig. 1a multiplied by an “inactivation” *r* gate representing the ‘down’ transitions (Clerx et al., 2019a; Mirams, 2023), so in the below designs when we refer to “Hodgkin-Huxley” (HH) it is this interpretation of the model we are using.

### 2.2 Wang model

The Wang et al. (1997) model can be written as:

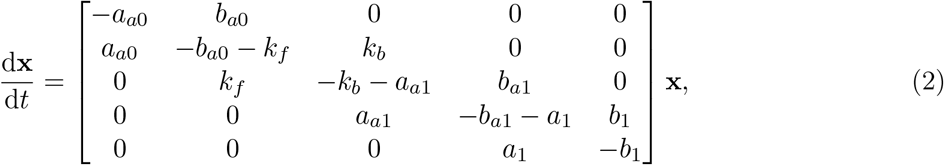

where

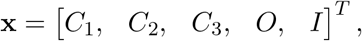

and

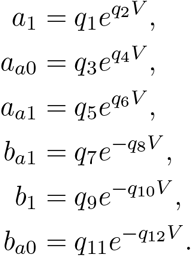

The default (room temperature) parameter values for both models are presented in Table 1. In practice we remove one state from the system and set it equal to “one minus the sum of the rest” to solve the ODE system, to improve numerical stability. All models are solved using a Python package Myokit (Clerx et al., 2016) using SUNDIALS CVODE (Hindmarsh et al., 2005).

**Table 1:**
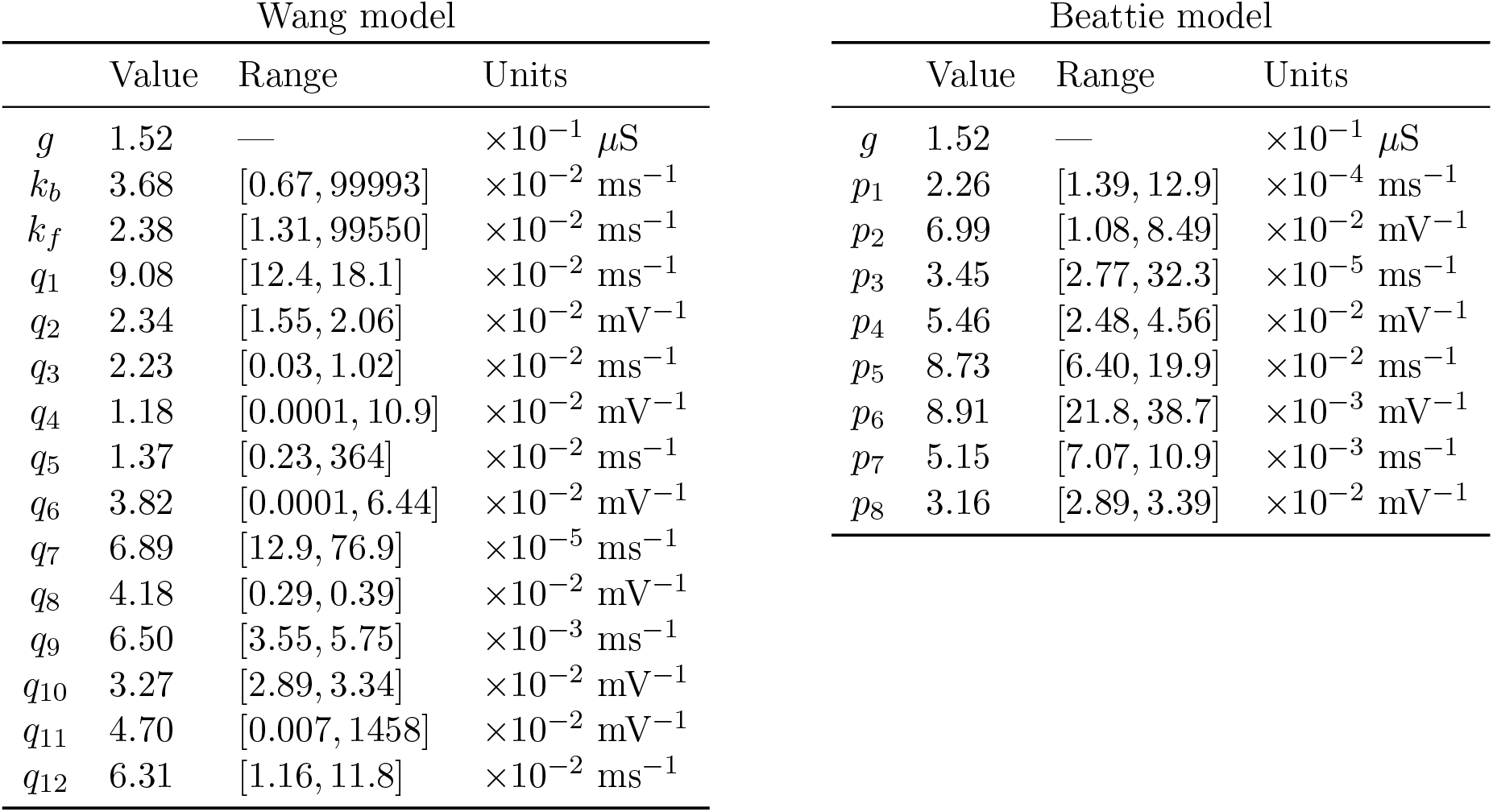
The default parameter sets we use for the Wang et al. (1997) and Beattie et al. (2018) models. The same maximal conductance (*g*) is used for both models. The column ‘Range’ indicates the parameter range obtained from real data fitting results based on protocols staircaseramp, sis, hh3step, and wang3step, which is used for global sensitivity-based designs.

| Wang model |  |  |  | Beattie model |  |  |  |
| --- | --- | --- | --- | --- | --- | --- | --- |
|  | Value | Range | Units |  | Value | Range | Units |
| $g$ | 1.52 | — | $\times 10^{-1} \mu\text{S}$ | $g$ | 1.52 | — | $\times 10^{-1} \mu\text{S}$ |
| $k_b$ | 3.68 | [0.67, 99993] | $\times 10^{-2} \text{ms}^{-1}$ | $p_1$ | 2.26 | [1.39, 12.9] | $\times 10^{-4} \text{ms}^{-1}$ |
| $k_f$ | 2.38 | [1.31, 99550] | $\times 10^{-2} \text{ms}^{-1}$ | $p_2$ | 6.99 | [1.08, 8.49] | $\times 10^{-2} \text{mV}^{-1}$ |
| $q_1$ | 9.08 | [12.4, 18.1] | $\times 10^{-2} \text{ms}^{-1}$ | $p_3$ | 3.45 | [2.77, 32.3] | $\times 10^{-5} \text{ms}^{-1}$ |
| $q_2$ | 2.34 | [1.55, 2.06] | $\times 10^{-2} \text{mV}^{-1}$ | $p_4$ | 5.46 | [2.48, 4.56] | $\times 10^{-2} \text{mV}^{-1}$ |
| $q_3$ | 2.23 | [0.03, 1.02] | $\times 10^{-2} \text{ms}^{-1}$ | $p_5$ | 8.73 | [6.40, 19.9] | $\times 10^{-2} \text{ms}^{-1}$ |
| $q_4$ | 1.18 | [0.0001, 10.9] | $\times 10^{-2} \text{mV}^{-1}$ | $p_6$ | 8.91 | [21.8, 38.7] | $\times 10^{-3} \text{mV}^{-1}$ |
| $q_5$ | 1.37 | [0.23, 364] | $\times 10^{-2} \text{ms}^{-1}$ | $p_7$ | 5.15 | [7.07, 10.9] | $\times 10^{-3} \text{ms}^{-1}$ |
| $q_6$ | 3.82 | [0.0001, 6.44] | $\times 10^{-2} \text{mV}^{-1}$ | $p_8$ | 3.16 | [2.89, 3.39] | $\times 10^{-2} \text{mV}^{-1}$ |
| $q_7$ | 6.89 | [12.9, 76.9] | $\times 10^{-5} \text{ms}^{-1}$ | | | | |
| $q_8$ | 4.18 | [0.29, 0.39] | $\times 10^{-2} \text{mV}^{-1}$ | | | | |
| $q_9$ | 6.50 | [3.55, 5.75] | $\times 10^{-3} \text{ms}^{-1}$ | | | | |
| $q_{10}$ | 3.27 | [2.89, 3.34] | $\times 10^{-2} \text{mV}^{-1}$ | | | | |
| $q_{11}$ | 4.70 | [0.007, 1458] | $\times 10^{-2} \text{mV}^{-1}$ | | | | |
| $q_{12}$ | 6.31 | [1.16, 11.8] | $\times 10^{-2} \text{ms}^{-1}$ | | | | |

## 3 Common protocol segments

As described in Mirams et al. (2024), all the protocols we have designed have a common start and end sections defined in Table 2. The purposes of these are

**Table 2:** Reproduced from Mirams et al. (2024). Details of the Start and End clamp sections for all designs. ‘*t*’ indicates the duration of the clamp section, and ‘*V* ‘ the relevant voltage(s) for this clamp. Where ‘Ramp’ is specified it is a linear ramp over time between the voltages shown, as opposed to a constant voltage clamp for a ‘Step’.

| Clamp # | Initial: for leak and conductance |  |  | End: reversal ramp sequence |  |  |
| --- | --- | --- | --- | --- | --- | --- |
| | Step/Ramp | $t$ (ms) | $V$ (mV) | Step/Ramp | $t$ (ms) | $V$ (mV) |
| 1 | Step | 250 | −80 | Step | 1000 | −80 |
| 2 | Step | 50 | −120 | Step | 500 | 40 |
| 3 | Ramp | 400 | −120 to −80 | Step | 10 | −70 |
| 4 | Step | 200 | −80 | Ramp | 100 | −70 to −110 |
| 5 | Step | 1000 | 40 | Step | 390 | −120 |
| 6 | Step | 500 | −120 | Step | 500 | −80 |
| 7 | Step | 1000 | −80 | — | — | — |

- Start — an ‘activation step’ to provoke a very large tail current and help with conductance estimation, as discussed in Beattie et al. (2018).
- End — a ‘reversal ramp’ to help assess whether the current is reversing at the expected Nernst potential, discussed in Lei et al. (2019b).
- both can also be used in quality control to check that these sections behave similarly over time when different protocols are applied to the same cell.

## 4 Manual protocol designs

The details of the protocols in this section are provided in Table 3.

**Table 3:** Details of the 5 protocols: staircase, sis, sisi, manualppx, and squarewave. All voltage values shown here are voltage steps to clamp to. These steps need to have the two ‘bookend’ sections added (see Table 2) which are identical for all designs.

| Clamp<br># | staircase |  | sis |  | sisi |  | manualppx |  | squarewave |  |
| --- | --- | --- | --- | --- | --- | --- | --- | --- | --- | --- |
|  | V (mV) | t (ms) | V (mV) | t (ms) | V (mV) | t (ms) | V (mV) | t (ms) | V (mV) | t (ms) |
| 1 | -40 | 500 | -40 | 500 | 40 | 500 | 60 | 200 | 60 | 24.9 |
| 2 | -60 | 500 | -60 | 500 | 0 | 500 | -60 | 200 | 40 | 25 |
| 3 | -20 | 500 | -20 | 500 | 20 | 500 | -100 | 200 | 60 | 25 |
| 4 | -40 | 500 | -40 | 500 | -20 | 500 | 40 | 500 | 40 | 25 |
| 5 | 0 | 500 | 0 | 500 | 0 | 500 | -90 | 200 | 60 | 25.1 |
| 6 | -20 | 500 | -20 | 500 | -40 | 500 | 30 | 500 | 40 | 9.9 |
| 7 | 20 | 500 | 20 | 500 | -20 | 500 | -80 | 200 | -12 | 15 |
| 8 | 0 | 500 | 0 | 500 | -60 | 500 | -100 | 200 | 8 | 25.1 |
| 9 | 40 | 500 | 40 | 225 | -40 | 225 | 20 | 200 | -12 | 24.9 |
| 10 | 20 | 500 | -80 | 50 | -80 | 50 | -40 | 1000 | 8 | 25 |
| 11 | 40 | 500 | -40 | 50 | -40 | 50 | 60 | 200 | -12 | 25.1 |
| 12 | 0 | 500 | -60 | 50 | -60 | 50 | 0 | 200 | 8 | 19.9 |
| 13 | 20 | 500 | -20 | 50 | -20 | 50 | -50 | 1000 | 60 | 5 |
| 14 | -20 | 500 | -40 | 50 | -40 | 50 | -10 | 100 | 40 | 25 |
| 15 | 0 | 500 | 0 | 50 | 0 | 50 | 10 | 100 | 60 | 25.1 |
| 16 | -40 | 500 | -20 | 50 | -20 | 50 | -20 | 100 | 40 | 25 |
| 17 | -20 | 500 | 20 | 50 | 20 | 50 | -80 | 300 | 60 | 25 |
| 18 | -60 | 500 | 0 | 50 | 0 | 50 | 0 | 100 | 40 | 24.9 |
| 19 | -40 | 500 | 40 | 50 | 40 | 50 | -20 | 100 | 60 | 5 |
| 20 | — | — | 20 | 50 | 20 | 50 | -100 | 200 | 8 | 20.1 |
| 21 |  |  | 40 | 50 | 40 | 50 | 40 | 300 | -12 | 24.9 |
| 22 |  |  | 0 | 50 | 0 | 50 | -60 | 100 | 8 | 25 |
| 23 |  |  | 20 | 50 | 20 | 50 | 0 | 100 | -12 | 25.1 |
| 24 |  |  | -20 | 50 | -20 | 50 | -10 | 100 | 8 | 24.9 |
| 25 |  |  | 0 | 50 | 0 | 50 | -20 | 100 | -12 | 15 |
| 26 |  |  | -40 | 50 | -40 | 50 | -30 | 100 | 40 | 10 |
| 27 |  |  | -20 | 50 | -20 | 50 | -40 | 100 | 60 | 25.1 |
| 28 |  |  | -60 | 50 | -60 | 50 | -80 | 100 | 40 | 24.9 |
| 29 |  |  | -40 | 50 | -40 | 50 | 30 | 100 | 60 | 25 |
| 30 |  |  | -80 | 50 | -80 | 50 | 60 | 100 | 40 | 25.1 |
| 31 |  |  | 40 | 225 | -40 | 225 | — | — | 60 | 24.9 |
| 32 |  |  | 0 | 500 | -60 | 500 |  |  | -12 | 25.1 |
| 33 |  |  | 20 | 500 | -20 | 500 |  |  | -100 | 25 |
| 34 |  |  | -20 | 500 | -40 | 500 |  |  | -120 | 25 |
| 35 |  |  | 0 | 500 | 0 | 500 |  |  | -100 | 25 |
| 36 |  |  | -40 | 500 | -20 | 500 |  |  | -120 | 24.9 |
| 37 |  |  | -20 | 500 | 20 | 500 |  |  | -100 | 10 |
| 38 |  |  | -60 | 500 | 0 | 500 |  |  | -48 | 15 |
| 39 |  |  | -40 | 500 | 40 | 500 |  |  | -68 | 25.1 |
| 40 |  |  | — | — | — | — |  |  | -48 | 25 |
| 41 |  |  |  |  |  |  |  |  | -68 | 24.9 |
| 42 |  |  |  |  |  |  |  |  | -48 | 25 |
| 43 |  |  |  |  |  |  |  |  | -68 | 20 |
| 44 |  |  |  |  |  |  |  |  | -120 | 5 |
| 45 |  |  |  |  |  |  |  |  | -100 | 25 |
| 46 |  |  |  |  |  |  |  |  | -120 | 25.1 |
| 47 |  |  |  |  |  |  |  |  | -100 | 25 |
| 48 |  |  |  |  |  |  |  |  | -120 | 24.9 |
| 49 |  |  |  |  |  |  |  |  | -100 | 25.1 |
| 50 |  |  |  |  |  |  |  |  | -120 | 4.9 |
| 51 |  |  |  |  |  |  |  |  | -68 | 20 |

### 4.1 Original staircase protocol

Figure 2 shows the original staircase protocol. It was manually designed to capture various dynamics of hERG (Lei et al., 2019b,a), which has been used and tested on the Nanion SyncroPatch384PE. We have been using it as a quality control of the full run of the experiments when designing the protocols in the rest of this report.

### 4.2 Staircase-in-staircase protocol

The original staircase protocol provided a good foundation and motivation for improving experimental designs for characterisation of ion channel kinetics in high-throughput machines. We attempted to further improve this manual design by enhancing the exploration of inactivation processes of hERG. The original staircase protocol involves only voltage steps of 500 ms, which may not be able to explore fully the fast dynamics of hERG inactivation processes. Therefore, a shorter step duration version (50 ms) of the full staircase protocol is introduced at the middle of the staircase protocol, termed the staircase-in-staircase (sis) protocol (Figure 3). We also explored the possibility of inverting the order of the staircase as shown in Figure 4 (sisi).

### 4.3 Phase-space filling protocol

The idea here is to have a protocol that fills up the phase-voltage space as much as possible. In brief, this design draws out the *a, r, V* three dimensional ‘phase-voltage space’ {[0, 1], [0, 1], [−120, 60]} for the Beattie et al. (2018) model and subdivides it into 6 compartments in each dimension, giving a total of *N* = 6^3^ = 216 boxes. Since the phase space defines all possible behaviours of a model, if a protocol forces the model to visit as many of these boxes as possible, then the observations should test model assumptions well and provide rich information to fit model parameters. We have published the rationale and details of the design process for these protocol separately in Mirams et al. (2024). Figure 5 shows a manually-tuned phase space filling protocol (manualppx); no objective function *per se*.

**Figure 5.**
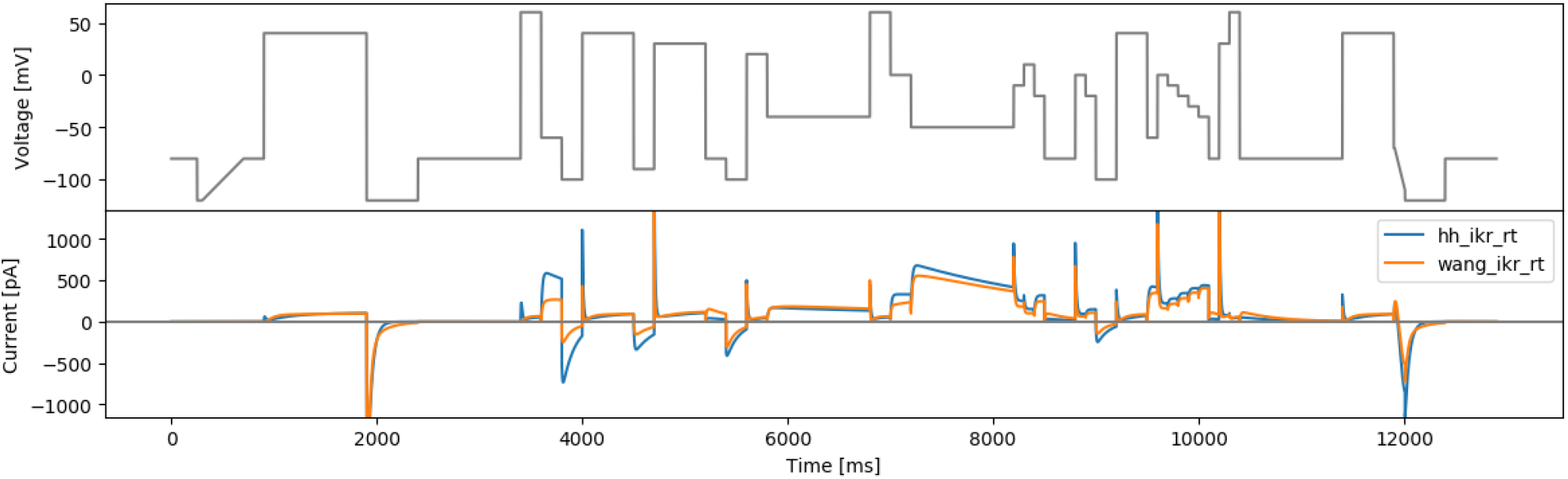
The manual phase space protocol (manualppx) and simulated currents from both models.

### 4.4 A square-wave conversion of the sinusoidal protocol

In this design, we aim to design protocols based on sums of square waves, as inspired by Beattie et al. (2018). Such a protocol consists of a combination of *N* square waves, where each square wave *i* is defined by amplitude *a*_*i*_, (angular) frequency *ω*_*i*_, and phase lag *ϕ*_*i*_. The protocol is defined by 3*N* parameters plus an extra parameter for an offset voltage, which can be expressed as:

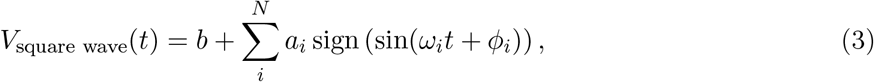

where the function sign(·) takes a value +1 if its argument is positive, −1 if negative, or 0 if the argument is 0.

A direct conversion of the sine waves in the Beattie et al. (2018) protocol is performed, with the same amplitudes and frequencies, to square waves. It is a combination of three square waves (*N* = 3) with *a*_1_ =54 mV, *a*_2_ =26 mV, *a*_3_ =10 mV, *ω*_1_ =0.007 ms^−1^, *ω*_2_ =0.037 ms^−1^, *ω*_3_ =0.19 ms^−1^, and *ϕ*_1_ = *ϕ*_2_ = *ϕ*_3_ = 0, and an offset of *b* =−30 mV. The resulting protocol is called ‘squarewave’ and is shown in Figure 6.

**Figure 6.**
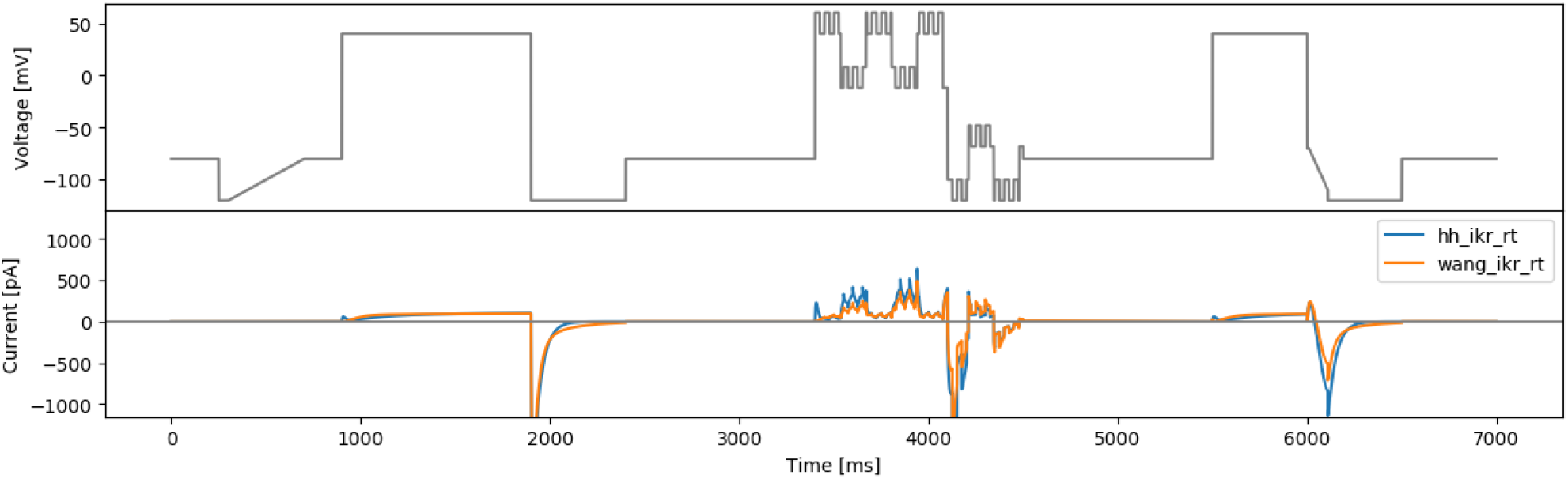
The square wave protocol of Beattie et al. (2018) (squarewave) and simulated currents from both models.

### 4.5 Long action potential protocol

As a final ‘manually-chosen’ design, we also propose a lumped action potential protocol for validation purposes, as shown in Figure 7. It consists of two action potential morphologies, an early after-depolarisation (EAD)-like action potential, and a delayed after-depolarisation (DAD)-like action potential. The details of this longap protocol are provided in Table 6.

**Figure 7.**
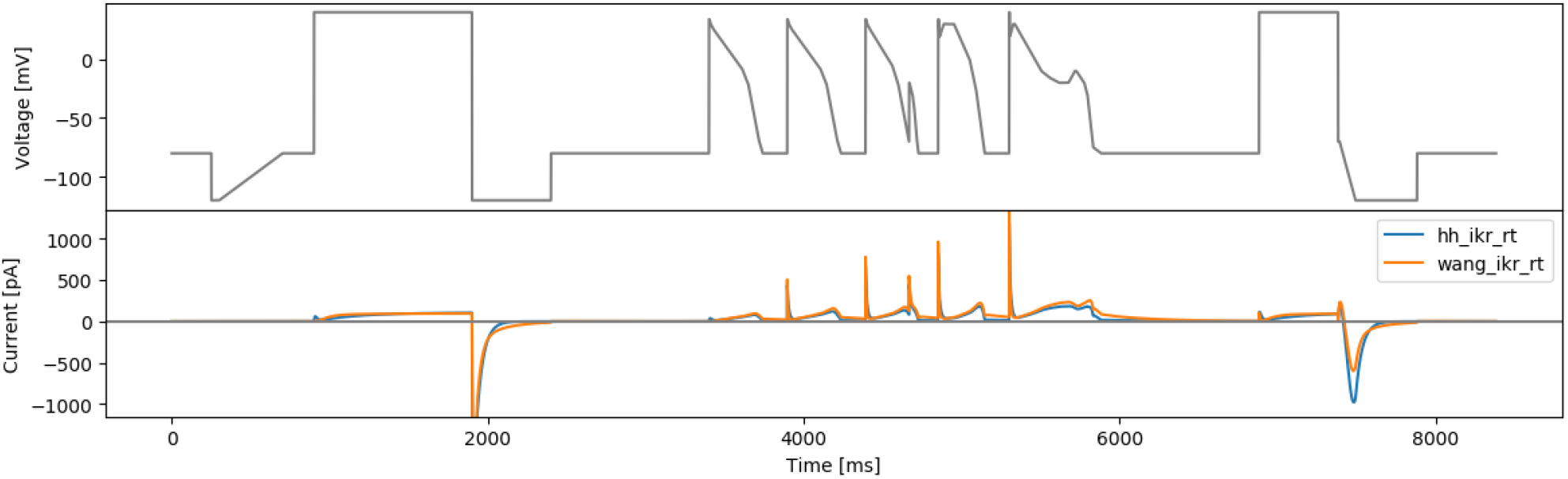
The lumped action potential protocol (longap) and simulated currents from both models.

## 5 Automated Iterative 3-step designs

Here we describe protocol design approaches that can be done objectively and automatically. With the same rationale as described in Mirams et al. (2024), we consider a protocol consists of 3*N* steps with *N* ∈ ℕ, and we split the protocol into *N* units with 3 consecutive voltage steps as a unit. For some designs, *N* is the number of model parameters, while for others, *N* is 17 to bring the total number of steps to 51 which is close to the 64 allowed by the Nanion SyncroPatch384PE when the start and end clamps are added (Table 2). For each unit *i*, we optimise the 3 voltage steps through an objective function *S*_*i*_, with each step defined by two parameters: voltage *V* and duration Δ*t*. Each objective function *S*_*i*_ (described in the sections below) aims to achieve a different purpose. We then iterate the process for all the objective functions *i* = 1, 2, …, *N*, resulting in a 3*N* steps protocol.

The optimisation was performed using a global optimisation scheme, covariance matrix adaptation evolution strategy (CMA-ES, Hansen, 2006) implemented via a Python package PINTS (Clerx et al., 2019b). All optimisation of the designs were repeated 10 times from different randomly varied initial starting points, and the best designs are presented here. Although we do not expect our design would reach the same global optimum as optimising all > 20 steps at once (Mirams et al., 2024), our results still show promising protocol designs. We also tried to perform fitting 6-steps-at-once in Mirams et al. (2024) and showed that both resulted in similar performance. Finally, the presented results are the optimised results rounded to the nearest one decimal place in millisecond and millivolt for practical implementation (Mirams et al., 2024).

The details of the protocols in this section are provided in Tables 4, 5, and 6.

**Table 4:** Details of the 5 protocols: hh3step, wang3step, hhsobol3step, wangsobol3step, and spacefill26. All voltage values shown here are voltage steps to clamp to. These steps need to have the two ‘bookend’ sections added (see Table 2) which are identical for all designs.

| Clamp<br># | hh3step |  | wang3step |  | hhsobol3step |  | wangsobol3step |  | spacefill26 |  |
| --- | --- | --- | --- | --- | --- | --- | --- | --- | --- | --- |
|  | V (mV) | t (ms) | V (mV) | t (ms) | V (mV) | t (ms) | V (mV) | t (ms) | V (mV) | t (ms) |
| 1 | -96.7 | 983 | 59.8 | 1000 | 60 | 1000 | -120 | 52.8 | 40 | 841 |
| 2 | 59.7 | 730 | 35.3 | 1000 | 60 | 1000 | 48.2 | 1000 | -63 | 773 |
| 3 | -50.9 | 266 | -47.6 | 216 | -69.4 | 54.9 | -46.1 | 1000 | -117.9 | 163 |
| 4 | 9.07 | 852 | -107 | 341 | -82.5 | 999 | 1.3 | 1000 | 59.8 | 174 |
| 5 | 45.8 | 621 | -89.9 | 641 | 10.8 | 955 | -4.03 | 1000 | 22.1 | 46 |
| 6 | -45.5 | 360 | -80.7 | 377 | -51.1 | 219 | -17.8 | 1000 | -97 | 214 |
| 7 | -120 | 999 | -119 | 998 | -81.8 | 999 | -97.7 | 50.4 | 32.9 | 409 |
| 8 | -120 | 1000 | -74.6 | 281 | 60 | 51.5 | -85 | 784 | -106.1 | 29 |
| 9 | -88 | 222 | -60.4 | 54.4 | -55.4 | 103 | -85 | 232 | 25.1 | 20 |
| 10 | 30.2 | 388 | 59.8 | 1000 | -80.2 | 488 | -85.2 | 1000 | -86.8 | 23 |
| 11 | 56.6 | 972 | 28.3 | 1000 | 60 | 1000 | -89.8 | 711 | 59.9 | 56 |
| 12 | -120 | 50.2 | -47.8 | 233 | -71.1 | 1000 | -120 | 1000 | -76.9 | 156 |
| 13 | 57.5 | 497 | -111 | 61.2 | 60 | 1000 | -82.4 | 195 | -6 | 20 |
| 14 | -120 | 1000 | -99.2 | 398 | 60 | 1000 | 41.6 | 1000 | -74.3 | 37 |
| 15 | -120 | 999 | -78.7 | 116 | -120 | 102 | -57 | 108 | -10.6 | 20 |
| 16 | -96 | 642 | -102 | 783 | 60 | 1000 | -84.3 | 548 | -75 | 164 |
| 17 | 59.8 | 806 | -66.6 | 219 | 60 | 1000 | 5.02 | 261 | 55 | 160 |
| 18 | -42.5 | 400 | 60 | 151 | -7.19 | 269 | 51 | 129 | -47.5 | 25 |
| 19 | 56 | 936 | -97.6 | 443 | -64.9 | 50 | -99.5 | 1000 | -7.8 | 38 |
| 20 | -4.8 | 55.6 | -97.6 | 784 | -47.3 | 75.1 | 2.12 | 1000 | -74.4 | 213 |
| 21 | 59.8 | 50 | -107 | 317 | -81.4 | 67.3 | -41.7 | 187 | -42.7 | 367 |
| 22 | -53.1 | 488 | -95.3 | 665 | 60 | 1000 | -85.2 | 999 | -52.5 | 483 |
| 23 | 59 | 989 | -119 | 616 | 60 | 1000 | 41.9 | 50.2 | -85 | 33 |
| 24 | -42.8 | 321 | -111 | 407 | -1.7 | 1000 | -85.1 | 650 | 5.5 | 20 |
| 25 | -77.9 | 753 | 60 | 1000 | -52.4 | 50 | -69.6 | 1000 | -105.5 | 27 |
| 26 | 46.5 | 911 | -120 | 50 | 60 | 1000 | -10.2 | 815 | -58.6 | 32 |
| 27 | -116 | 54.3 | -120 | 50 | -54.1 | 50 | -71.5 | 1000 | -114.2 | 20 |
| 28 | — | — | 59.6 | 1000 | — | — | 20.3 | 1000 | 14.1 | 108 |
| 29 |  |  | 30.5 | 1000 |  |  | 46.8 | 797 | -90.5 | 20 |
| 30 |  |  | -39.7 | 297 |  |  | -120 | 663 | -49.1 | 20 |
| 31 |  |  | -120 | 725 |  |  | -8.4 | 128 | 59.9 | 103 |
| 32 |  |  | -106 | 225 |  |  | 36.1 | 374 | -101.7 | 20 |
| 33 |  |  | -108 | 568 |  |  | 53.8 | 999 | 15.1 | 20 |
| 34 |  |  | 59.6 | 1000 |  |  | -85 | 949 | -87.8 | 61 |
| 35 |  |  | 31.3 | 999 |  |  | -84.9 | 423 | 15.4 | 272 |
| 36 |  |  | -41.9 | 187 |  |  | -111 | 129 | -114 | 169 |
| 37 |  |  | 60 | 1000 |  |  | -120 | 1000 | 34.7 | 892 |
| 38 |  |  | 60 | 1000 |  |  | 22 | 198 | -83.5 | 87 |
| 39 |  |  | 60 | 1000 |  |  | -88.9 | 1000 | 46.6 | 444 |
| 40 |  |  | -66.1 | 727 |  |  | -84.6 | 869 | -100.2 | 23 |
| 41 |  |  | -120 | 931 |  |  | 33.6 | 50 | -3.3 | 23 |
| 42 |  |  | 0 | 50 |  |  | 27.3 | 99.3 | 21 | 26 |
| 43 |  |  | 60 | 159 |  |  | -120 | 50.3 | -55.8 | 421 |
| 44 |  |  | -120 | 1000 |  |  | -85 | 107 | -95.3 | 29 |
| 45 |  |  | -55.2 | 1000 |  |  | -85 | 60.7 | -8.4 | 32 |
| 46 |  |  | — | — |  |  | — | — | -101.6 | 33 |
| 47 |  |  |  |  |  |  |  |  | -20.7 | 20 |
| 48 |  |  |  |  |  |  |  |  | -64.9 | 20 |
| 49 |  |  |  |  |  |  |  |  | 50.5 | 585 |
| 50 |  |  |  |  |  |  |  |  | -97.4 | 115 |
| 51 |  |  |  |  |  |  |  |  | 3.7 | 658 |

**Table 5:** Details of the 5 protocols: spacefill10, spacefill19, hhbrute3gstep, wangbrute3gstep, and rvotmaxdiff. All voltage values shown here are voltage steps to clamp to. These steps need to have the two ‘bookend’ sections added (see Table 2) which are identical for all designs.

| Clamp<br># | spacefill10 |  | spacefill19 |  | hhbrute3gstep |  | wangbrute3gstep |  | rvotmaxdiff |  |
| --- | --- | --- | --- | --- | --- | --- | --- | --- | --- | --- |
|  | V (mV) | t (ms) | V (mV) | t (ms) | V (mV) | t (ms) | V (mV) | t (ms) | V (mV) | t (ms) |
| 1 | 50.5 | 336 | 60 | 142 | 37.7 | 795 | 60 | 837 | 19 | 500 |
| 2 | -97.3 | 89 | -69.5 | 844 | -120 | 261 | -43.7 | 506 | -32 | 50 |
| 3 | -12.7 | 20 | -106.3 | 58 | -36.6 | 735 | -120 | 892 | -31 | 50 |
| 4 | -88.5 | 67 | -11.6 | 33 | 41 | 231 | -71.3 | 1000 | -49 | 50 |
| 5 | 18.8 | 804 | 48.8 | 584 | -45.3 | 815 | -117 | 50 | 5 | 50 |
| 6 | -114.3 | 166 | -97 | 689 | -65.4 | 50.2 | -114 | 50 | 54 | 439 |
| 7 | 59 | 149 | 33.6 | 752 | -120 | 530 | 57.9 | 169 | 22 | 499 |
| 8 | -60.5 | 438 | -79.1 | 398 | 60 | 459 | -33.4 | 617 | 22 | 500 |
| 9 | -97.5 | 120 | -49.8 | 257 | -120 | 714 | -116 | 757 | 19 | 145 |
| 10 | 57.5 | 144 | 50.5 | 99 | -19 | 1000 | -51.9 | 50 | -26 | 89 |
| 11 | -52.4 | 496 | -104.2 | 32 | 20.2 | 485 | 41.7 | 50 | -66 | 50 |
| 12 | -75 | 465 | -33 | 35 | -64.1 | 1000 | 57.2 | 50.4 | -85 | 50 |
| 13 | 34.7 | 711 | -106.3 | 62 | 50.4 | 947 | 12.8 | 446 | 7 | 50 |
| 14 | -113.5 | 31 | 11.5 | 228 | -34.1 | 362 | -28.6 | 358 | 43 | 121 |
| 15 | -9.7 | 299 | -79.6 | 153 | -36.5 | 991 | -55.2 | 746 | -17 | 53 |
| 16 | -70 | 33 | 58.2 | 594 | -47.3 | 1000 | -15.6 | 1000 | -13 | 50 |
| 17 | 12.2 | 79 | -71.3 | 462 | -30.8 | 1000 | -82.6 | 50 | 9 | 500 |
| 18 | -98.1 | 21 | -24.4 | 110 | -72.9 | 50.1 | -94.9 | 152 | -95 | 50 |
| 19 | 45.5 | 168 | 17.1 | 617 | 58.8 | 650 | -48.9 | 339 | -16 | 500 |
| 20 | -85.9 | 59 | -96.7 | 38 | -120 | 471 | 60 | 293 | -48 | 153 |
| 21 | 33.7 | 25 | 59.1 | 720 | -41.7 | 762 | -120 | 76.3 | -13 | 500 |
| 22 | -97.5 | 76 | -47.8 | 351 | -47 | 1000 | 10.8 | 363 | -59 | 50 |
| 23 | -42.7 | 32 | -98.5 | 151 | 35.4 | 50 | -27.8 | 50.5 | -97 | 50 |
| 24 | -109.7 | 21 | -28.1 | 457 | 8.1 | 50 | -43.6 | 1000 | 48 | 460 |
| 25 | 0.3 | 177 | 58.8 | 96 | 50.8 | 914 | 60 | 986 | 48 | 52 |
| 26 | -86.8 | 144 | -41.3 | 336 | -32.1 | 376 | -120 | 228 | 27 | 50 |
| 27 | -23.3 | 455 | -56.1 | 526 | -120 | 251 | 60 | 672 | -8 | 50 |
| 28 | -106.3 | 33 | 58.4 | 144 | -29.2 | 50 | 44.6 | 50 | -64 | 50 |
| 29 | 54.6 | 20 | -99.3 | 31 | 1.81 | 1000 | 49.8 | 50 | -90 | 50 |
| 30 | -60 | 169 | 59.8 | 382 | -30.1 | 1000 | -117 | 62.2 | 23 | 500 |
| 31 | 59.9 | 153 | -28 | 886 | -46.8 | 576 | 60 | 448 | — | — |
| 32 | -74.2 | 29 | -119.4 | 20 | 46.4 | 905 | -44.1 | 817 |  |  |
| 33 | 5.3 | 20 | -16.4 | 221 | -34.9 | 783 | -120 | 561 |  |  |
| 34 | -29.5 | 933 | -106.3 | 58 | -17.8 | 1000 | 9.6 | 50 |  |  |
| 35 | -105.9 | 35 | 54.5 | 586 | -0.1 | 1000 | 28.6 | 50 |  |  |
| 36 | 38 | 29 | -107.9 | 146 | 15.3 | 50 | 43 | 50.4 |  |  |
| 37 | -91.2 | 80 | 59 | 123 | 50.4 | 913 | 60 | 153 |  |  |
| 38 | -19 | 493 | -101 | 21 | -34.9 | 835 | -120 | 957 |  |  |
| 39 | -115.6 | 1007 | 37.2 | 20 | -38.9 | 818 | 60 | 206 |  |  |
| 40 | 59.9 | 218 | -102.3 | 46 | -116 | 50.5 | 32.1 | 50 |  |  |
| 41 | -99.5 | 54 | 37.7 | 182 | 50.5 | 115 | -7.3 | 50.5 |  |  |
| 42 | -42.1 | 799 | -27.8 | 849 | -98.1 | 324 | 1.1 | 50 |  |  |
| 43 | -101.5 | 105 | -43.9 | 44 | 60 | 512 | 58.9 | 200 |  |  |
| 44 | 14.5 | 36 | -93.5 | 37 | -120 | 980 | -45 | 947 |  |  |
| 45 | 33.7 | 754 | 16.7 | 107 | -37.1 | 98.9 | -120 | 105 |  |  |
| 46 | 56.3 | 45 | -42.7 | 179 | — | — | — | — |  |  |
| 47 | -75.8 | 25 | -97.3 | 102 |  |  |  |  |  |  |
| 48 | 28.7 | 26 | -8 | 250 |  |  |  |  |  |  |
| 49 | -20.5 | 364 | 26.4 | 93 |  |  |  |  |  |  |
| 50 | -98.9 | 26 | -101.3 | 20 |  |  |  |  |  |  |
| 51 | 13.1 | 21 | 26.8 | 27 |  |  |  |  |  |  |

**Table 6:** Details of the 3 protocols: rtovmaxdiff, maxdiff, and longap. All voltage values for protocols rtovmaxdiff and maxdiff are voltage steps to clamp to. Protocol longap also indicates with ‘Ramp’ or ‘Step’; ‘Ramp’ is specified it is a linear ramp over time between the voltages shown, as opposed to a constant voltage clamp for a ‘Step’. These steps need to have the two ‘bookend’ sections added (see Table 2) which are identical for all designs.

| Clamp<br># | rtovmaxdiff |  | maxdiff |  | longap |  |  |
| --- | --- | --- | --- | --- | --- | --- | --- |
|  | V (mV) | t (ms) | V (mV) | t (ms) | Step/Ramp | V (mV) | t (ms) |
| 1 | 60 | 167 | -120 | 12.5 | Step | 34 | 3 |
| 2 | -65 | 516 | 60 | 12.5 | Ramp | 30 | 8 |
| 3 | -49 | 861 | -120 | 12.5 | Ramp | 26 | 15.2 |
| 4 | -100 | 587 | 60 | 12.5 | Ramp | -8 | 183.6 |
| 5 | 46 | 658 | -120 | 12.5 | Ramp | -21 | 39 |
| 6 | -60 | 446 | 60 | 12.5 | Ramp | -68 | 65.8 |
| 7 | 60 | 150 | -120 | 12.5 | Ramp | -80 | 25.2 |
| 8 | 4 | 185 | 60 | 12.5 | Step | -80 | 155.6 |
| 9 | -100 | 208 | -120 | 12.5 | Step | 34 | 3 |
| 10 | -74 | 935 | 60 | 12.5 | Ramp | 30 | 8 |
| 11 | 42 | 742 | -120 | 12.5 | Ramp | 26 | 15.2 |
| 12 | 29 | 751 | 60 | 12.5 | Ramp | -8 | 183.6 |
| 13 | 60 | 986 | -120 | 12.5 | Ramp | -21 | 39 |
| 14 | -100 | 866 | 60 | 12.5 | Ramp | -68 | 65.8 |
| 15 | -2 | 797 | -120 | 12.5 | Ramp | -80 | 25.2 |
| 16 | 60 | 177 | 60 | 12.5 | Step | -80 | 155.6 |
| 17 | -84 | 79 | -120 | 12.5 | Step | 34 | 3 |
| 18 | 60 | 943 | 60 | 12.5 | Ramp | 30 | 8 |
| 19 | 60 | 494 | -120 | 12.5 | Ramp | 26 | 15.2 |
| 20 | 32 | 666 | 60 | 12.5 | Ramp | -5 | 142.6 |
| 21 | 37 | 73 | -120 | 12.5 | Ramp | -21 | 38.4 |
| 22 | -100 | 380 | 60 | 12.5 | Ramp | -70 | 68.6 |
| 23 | 60 | 474 | -120 | 12.5 | Step | -20 | 2 |
| 24 | 12 | 101 | 60 | 12.5 | Ramp | -30 | 20 |
| 25 | -1 | 904 | -120 | 12.5 | Ramp | -40 | 10 |
| 26 | 60 | 162 | 60 | 12.5 | Ramp | -65 | 15 |
| 27 | 9 | 989 | -120 | 12.5 | Ramp | -80 | 12 |
| 28 | 60 | 323 | 60 | 12.5 | Step | -80 | 125 |
| 29 | 2 | 444 | -120 | 12.5 | Step | 34 | 3 |
| 30 | -50 | 492 | 60 | 12.5 | Ramp | 19 | 6 |
| 31 | — | — | -120 | 12.5 | Ramp | 30 | 26.4 |
| 32 |  |  | 60 | 12.5 | Step | 30 | 65 |
| 33 |  |  | -120 | 12.5 | Ramp | 0 | 99 |
| 34 |  |  | 60 | 12.5 | Ramp | -25 | 40 |
| 35 |  |  | -120 | 12.5 | Ramp | -80 | 55 |
| 36 |  |  | 60 | 12.5 | Step | -80 | 155 |
| 37 |  |  | -120 | 12.5 | Step | 40 | 3 |
| 38 |  |  | 60 | 12.5 | Step | 20 | 3 |
| 39 |  |  | -120 | 12.5 | Ramp | 30 | 20 |
| 40 |  |  | 60 | 12.5 | Step | 30 | 10 |
| 41 |  |  | -120 | 12.5 | Ramp | -10 | 168 |
| 42 |  |  | 60 | 12.5 | Ramp | -15.5 | 50.6 |
| 43 |  |  | -120 | 12.5 | Ramp | -20 | 61.2 |
| 44 |  |  | 60 | 12.5 | Step | -20 | 60 |
| 45 |  |  | -120 | 12.5 | Ramp | -10 | 40 |
| 46 |  |  | 60 | 12.5 | Step | -10 | 10 |
| 47 |  |  | -120 | 12.5 | Ramp | -20 | 50 |
| 48 |  |  | 60 | 12.5 | Ramp | -30 | 20 |
| 49 |  |  | -120 | 12.5 | Ramp | -75 | 36 |
| 50 |  |  | 60 | 12.5 | Ramp | -80 | 50 |

### 5.1 Sensitivity-based designs

#### 5.1.1 Maximising approximated local sensitivity

For an ion channel current model *I* with *N* parameters *p*_1_, *p*_2_, …, *p*_*N*_, we define an objective function for each 3-step unit *i* that maximises the absolute value of the elasticity 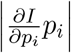 of the model output *I* with respect to the parameter *p*_*i*_ while minimising all the absolute value of elasticity of the rest of the parameters. This objective function can be mathematically expressed as

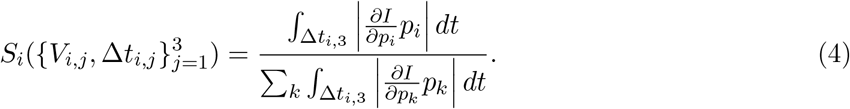

The sensitivity was calculated using a first-order central difference scheme with *δp*_*i*_ being 0.1 %×*p*_*i*_. Note that the integration is only over the last step of the 3 steps, the idea is to allow the first two steps to vary as much as it would need to be to maximise the approximated local sensitivity across the third step (it is fine if there is low sensitivity because of e.g. full inactivation in the first two steps). This has been repeated for both models and the results are shown in Figures 8 and 9.

**Figure 8.**
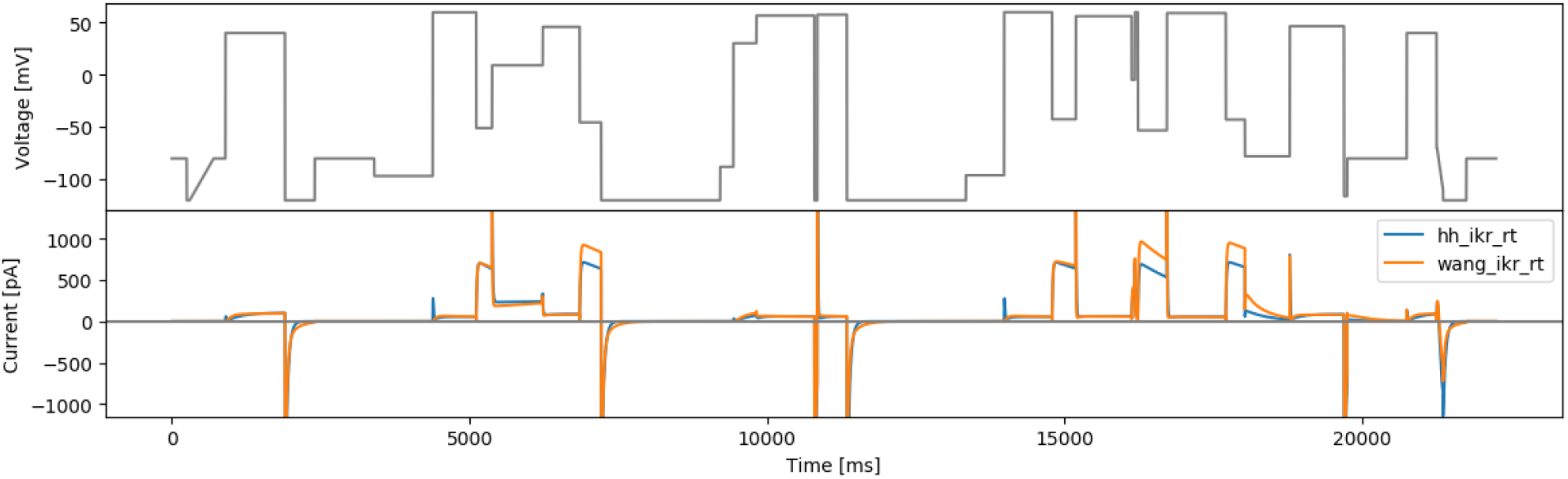
The 3-step local sensitivity protocol based on the Hodgkin-Huxley model (hh3step) and simulated currents from both models.

**Figure 9.**
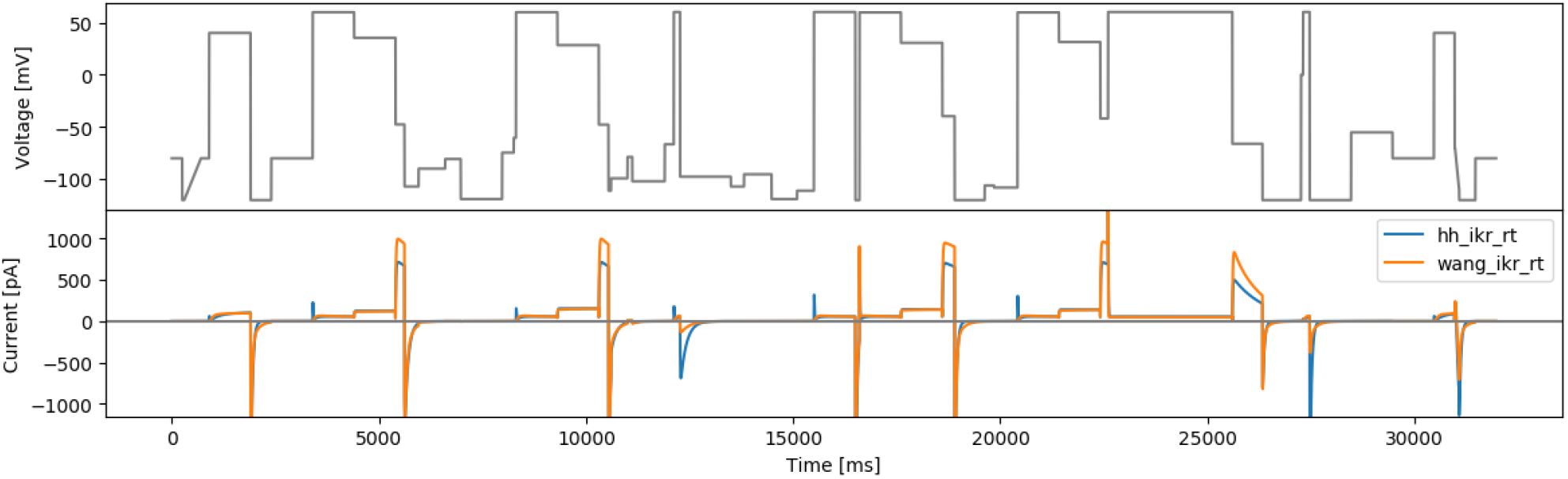
The 3-step local sensitivity protocol based on the Wang model (wang3step) and simulated currents from both models.

#### 5.1.2 Maximising Sobol sensitivity

Instead of the local sensitivity, we can also replace it with the first-order Sobol global sensitivity indices, given by

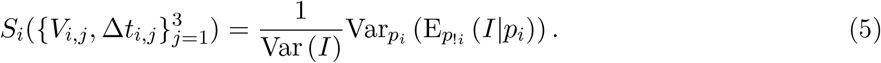

Here the *p*_!*i*_ notation denotes the set of all parameters except *p*_*i*_. This has been repeated for the two models described in Section 2. The parameter range (Table 1) was taken from previous real data fits to staircaseramp, sis, hh3step and wang3step, using the approach from Lei et al. (2019b) without accounting for experimental error (Lei et al., 2020a,b).

To calculate Sobol sensitivities we used a modified version of the SA-lib library (Herman and Usher, 2017), to enable easier calculation of sensitivities over time series, which is included in our repository (see Data Availability). The results are shown in Figure 10 and 11.

**Figure 10.**
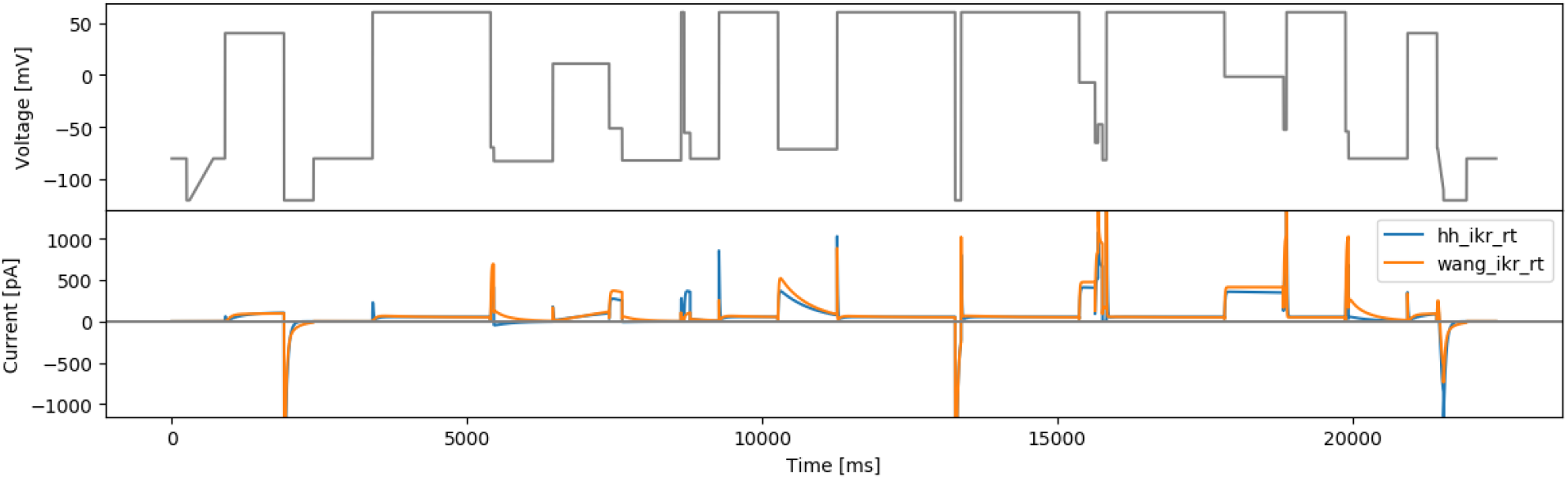
The 3-step Sobol sensitivity protocol based on the Hodgkin-Huxley model (hhsobol3step) and simulated currents from both models.

**Figure 11.**
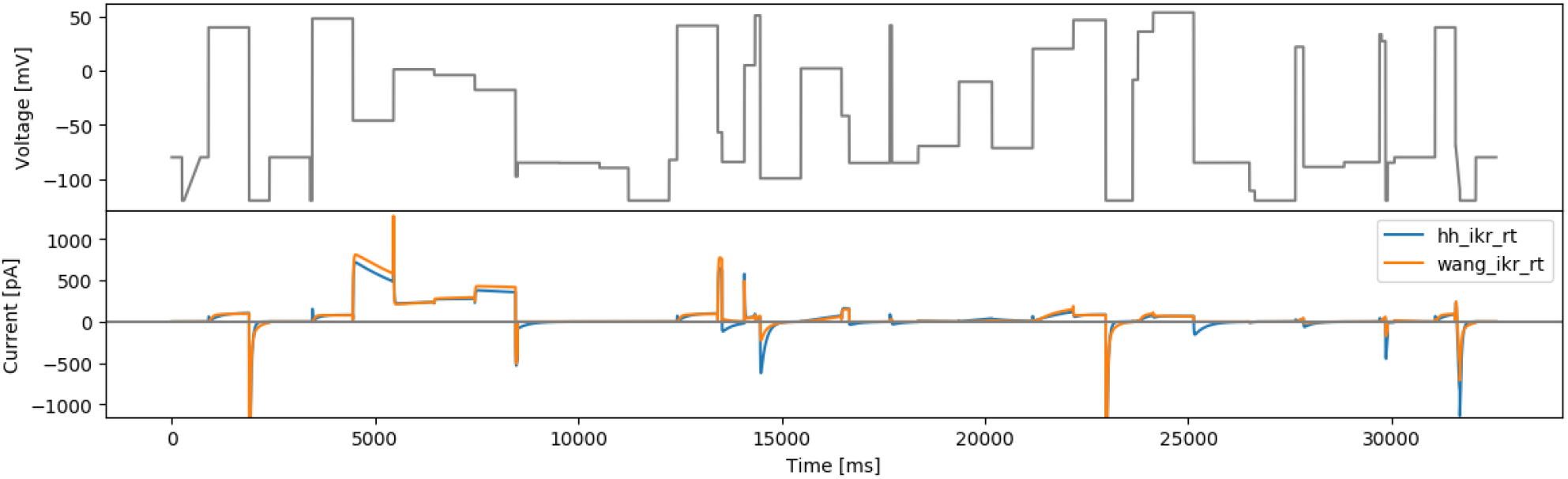
The 3-step Sobol sensitivity protocol based on the Wang model (wangsobol3step) and simulated currents from both models.

### 5.2 Phase-voltage space filling designs

For details of this approach, see Mirams et al. (2024). Briefly, an objective function tries to maximise the amount of new boxes that are visited by a model’s trajectory for each new iterative ‘3 step’ set of pulses (as described in Section 4.3) repeating sequentially until we have 17 sets of 3 steps. This approach has a stochastic optimisation step, and produces some protocols that appear to be challenging and information rich, where we appear to have a reasonable amount of current and interesting dynamics. After 30 optimisation runs with different random seeds and initial guesses, we selected the following 3 best protocols based on slightly different criteria:

- Figure 12 — Number 26: the best spcae-filling objective function score (Mirams et al., 2024).
- Figure 13 — Number 10: the best score for the RMSD value between the two models described in Section 2 as in Section 5.3.2.
- Figure 14 — Number 19: the best brute-force sampling score (Section 5.3.1) for the Beattie et al. (2018) model.

**Figure 12.**
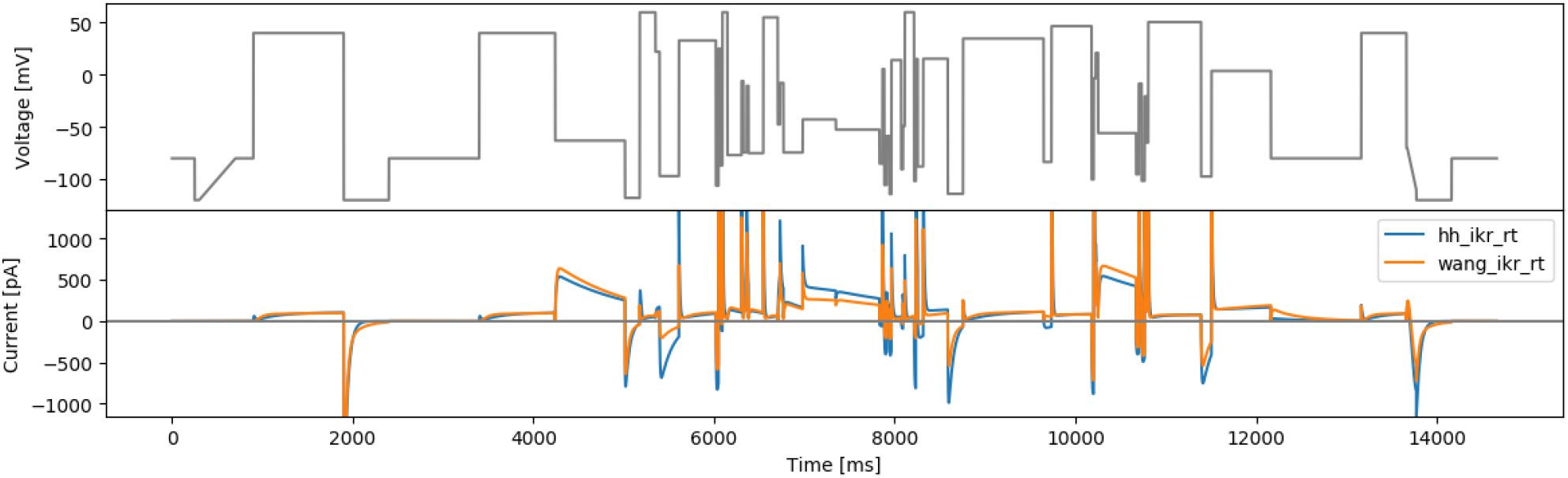
The first phase-voltage space protocol (spacefill26) and simulated currents from both models.

**Figure 13.**
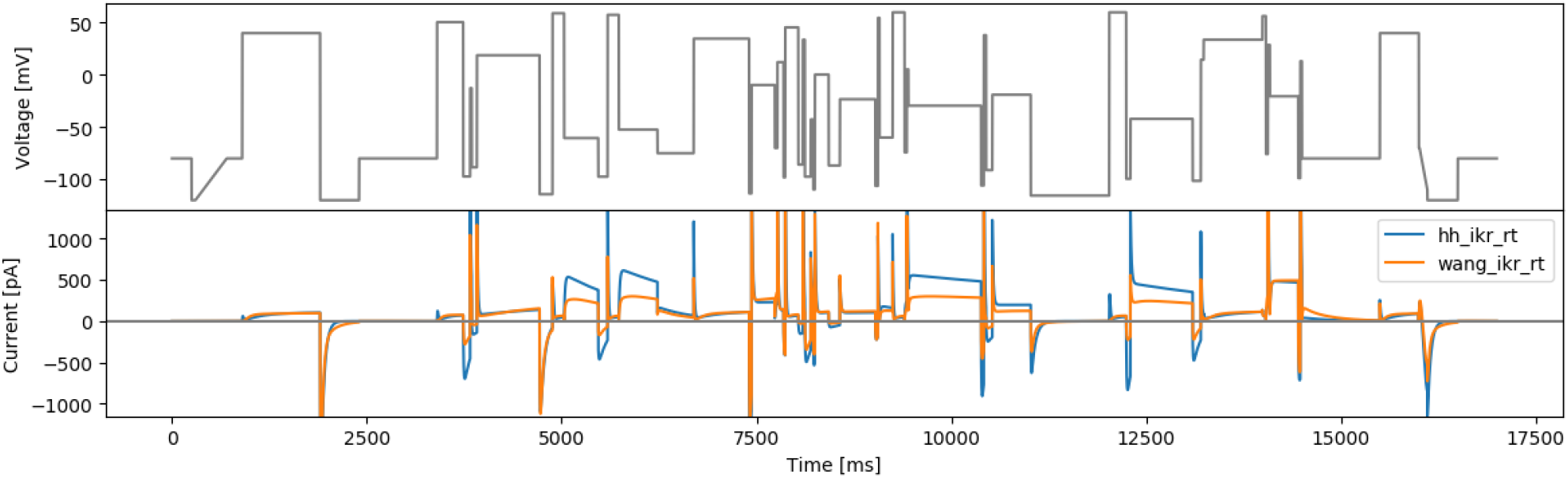
The second phase-voltage space protocol (spacefill10) and simulated currents from both models.

**Figure 14.**
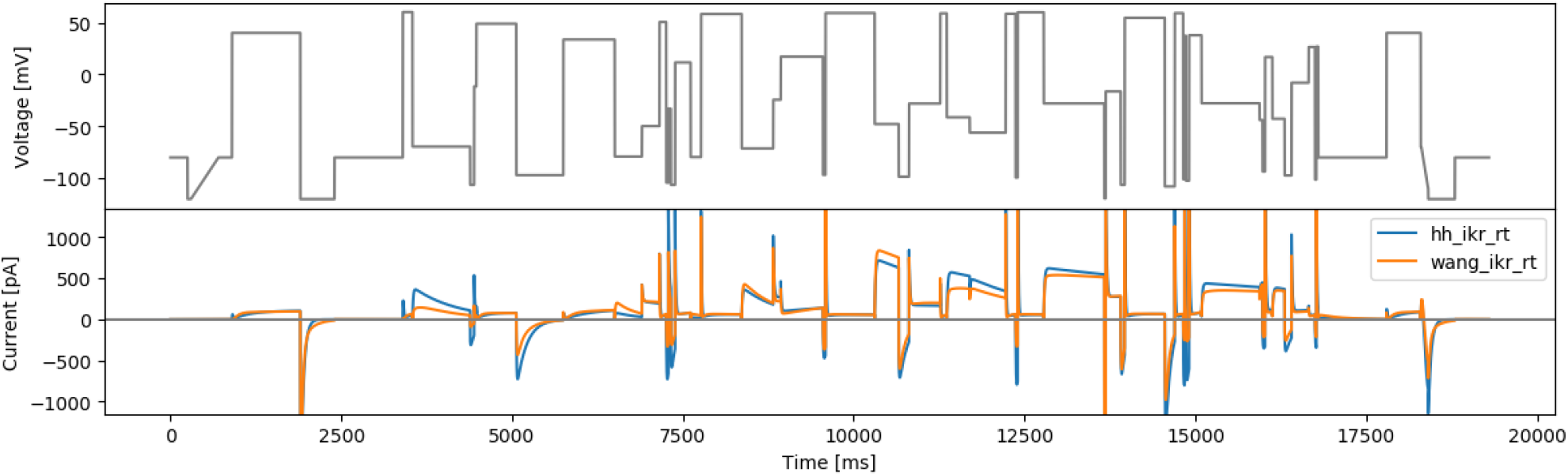
The third phase-voltage space protocol (spacefill19) and simulated currents from both models.

All three protocols visit between 126–132 (58–61%) of the available 216 ‘boxes’ in phase-voltage space. Note that this is a lower percentage than the protocols in Mirams et al. (2024) primarily due to 1 ms time samples being used in the 2019 optimisations presented here (see Discussion of Mirams et al. (2024)) along with extra initial guesses now being used in the Mirams et al. (2024) optimisation procedure to gain slightly higher coverage of the space.

### 5.3 Gibbs designs

We use the 3-step approach discussed above, but the difference here is that instead of defining each step by two parameters (voltage *V* and duration Δ*t*), for each 3-step section we optimise only one of these parameters (either *V* or Δ*t*) while randomly picking the other from a uniform distribution. This halves the number of parameters that are inferred to just 3 per 3-step section. However, since we have only the same objective function, all units would return the same optimum (or a few if multi-modal but very limited) which is not desired. Therefore we introduce some stochasticity to the protocol by randomly choosing one of the step parameters and optimising only the other one.

#### 5.3.1 Maximising model output differences: a brute-force sampling approach

The approach taken in this design is similar to a global sensitivity analysis. For a given model *I*, we start with randomly picking *M* (ideally ∼ 1000s but practically ∼ 100s of) parameters from model parameter prior, then the objective function to be optimised is the sum of the root mean square deviation (RMSD) values between the model outputs from all combinations of the sampled parameter pairs. The model parameter prior could be an a-priori distribution of the parameters (for example those used in Beattie et al., 2018; Lei et al., 2019b), or based on previous fitting results (see below). The objective function for a 3-step unit *i* can be expressed as

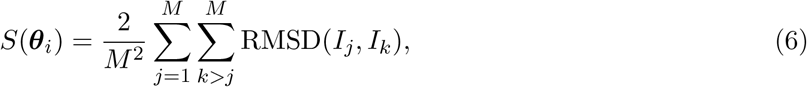

where RMSD(*x, y*) denotes the RMSD between *x* and *y*, and *I*_*j*_, *I*_*k*_ are the model output for the *M* parameter samples. We choose 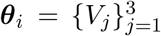 with Δ*t*_*j*_ ∼ Uniform(50, 1000) ms for odd *i*, and 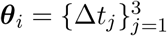 with *V*_*j*_ ∼ Uniform(−120, 60) mV for even *i*.

This has been repeated for the two models described in Section 2, and the parameter range (prior distribution) was taken from the maximum range defined by previous real data fits to staircsaeramp, sis, hh3step and wang3step, as provided in Table 1. The results are shown in Figure 15 and 16.

**Figure 15.**
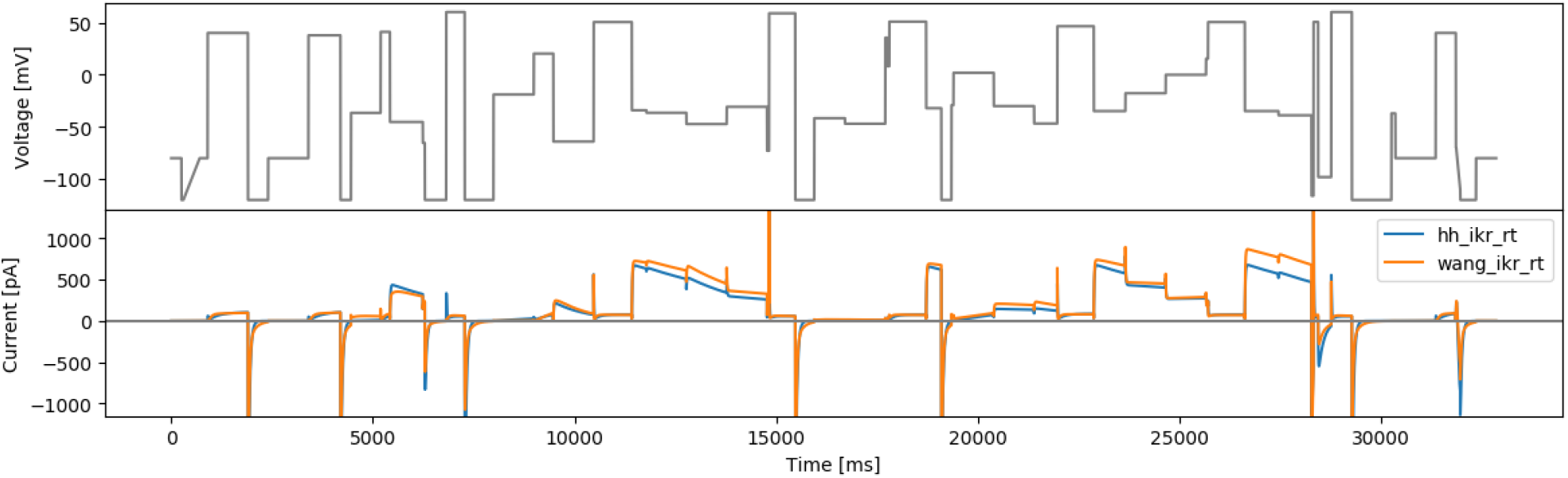
The brute-force sampling protocol based on the Hodgkin-Huxley model (hhbrute3gstep) and simulated currents from both models.

**Figure 16.**
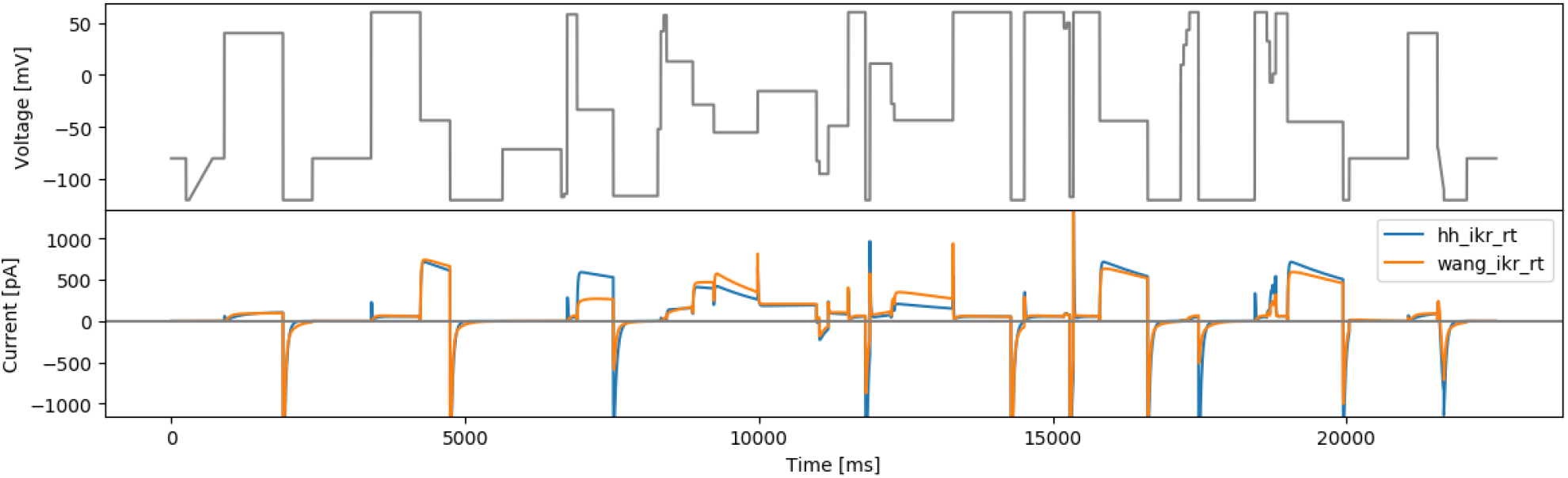
The brute-force sampling protocol based on the Wang model (wangbrute3gstep) and simulated currents from both models.

#### 5.3.2 Maximising differences between two models

Unlike the previously defined approaches, where only one model was involved, this proposed approach aims to distinguish between two candidate models. The objective function is defined as the RMSD value between two model currents, with a given set of model parameters (Table 1), so it is still a ‘local’ design with respect to model parameters. One protocol randomly picks time parameters for each 3-step unit, and optimises voltages (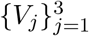 with Δ*t*_*j*_ ∼ Uniform(50, 500) ms and is termed ‘rtovmaxdiff’); and the other method randomly picks voltages and optimises the step durations 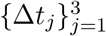 with *V*_*j*_ ∼ Uniform(−120, 60) mV, and is known as ‘rvotmaxdiff’. Applying this approach to the two models described in Section 2 results in Figures 17 and 18.

**Figure 17.**
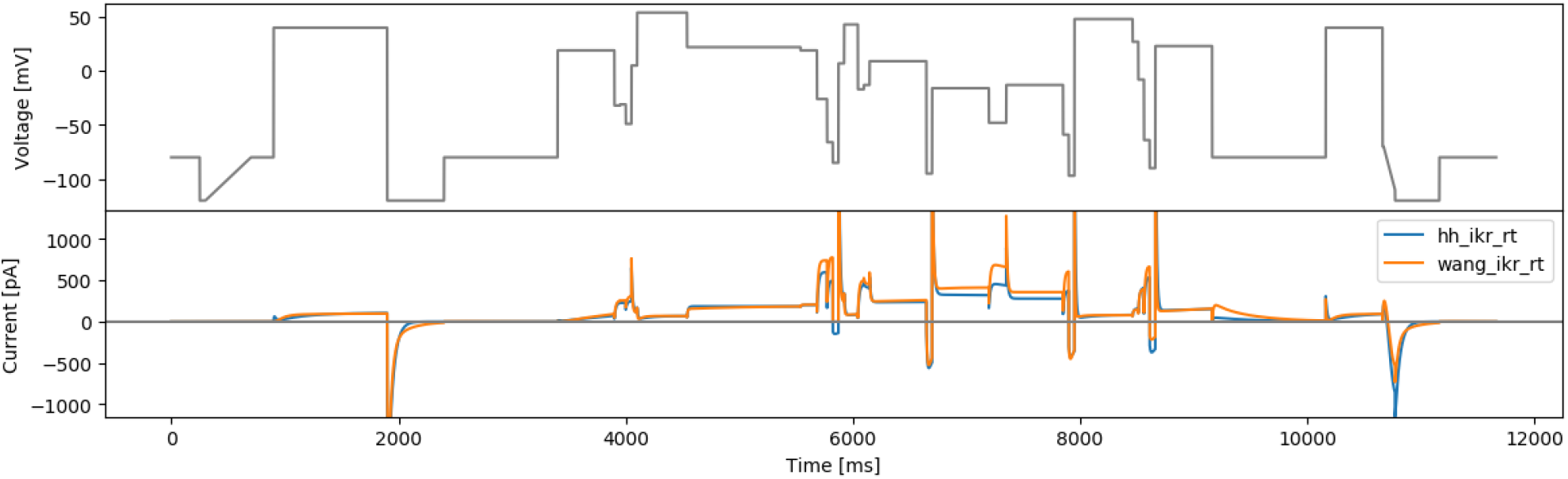
Protocol rvotmaxdiff and simulated currents from both models.

**Figure 18.**
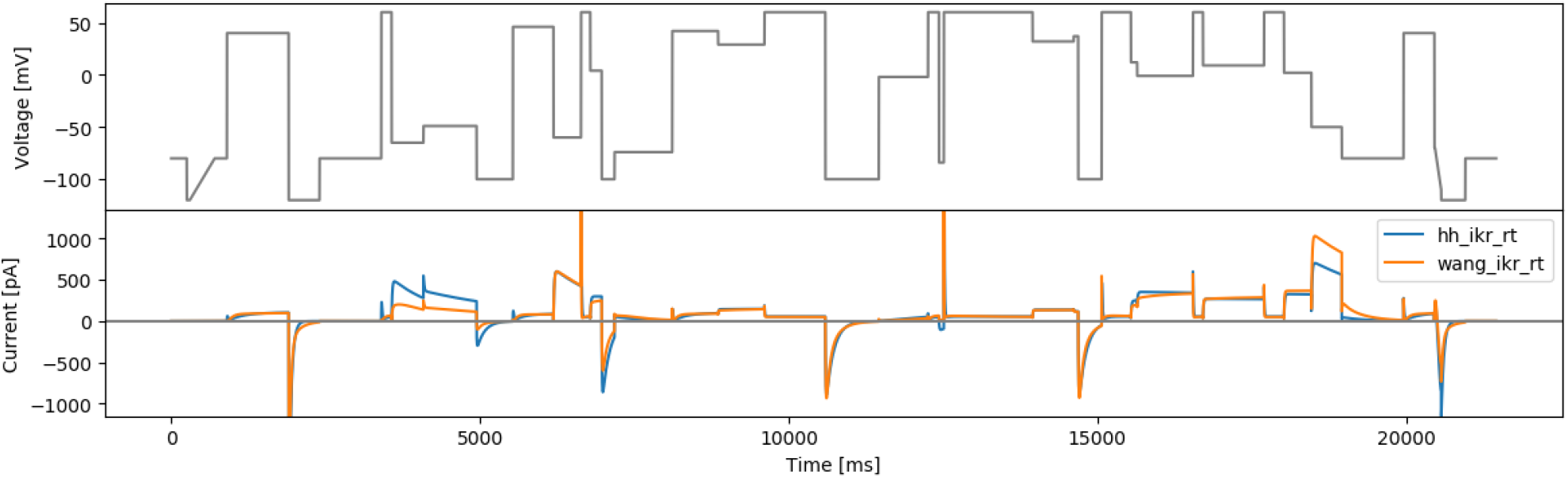
Protocol rtovmaxdiff and simulated currents from both models.

## 6 Automated square waves

Following the same argument as in Section 5.3.2, this design maximises the differences between two candidate models to aid model selection. Here we use *N* = 3 (as per Beattie et al., 2018) which gives 9 parameters in total (see Equation (3)), with a fixed offset voltage of −30 mV. The square wave parameters are optimised based on an objective function that maximises the RMSD value between two model outputs. Similar to Section 5.3.2, the two models have a set of predefined model parameters, so it is still a ‘local’ model parameter method.

This approach was applied to the two models described in Section 2 using the original model parameters. The resulting protocol (Figure 19) exhibits extremely high frequency and high amplitude (hitting the boundaries of the protocol parameters) behaviour. We believe these rapid changes of voltage tends to maximise the two model outputs, which is similar to the ‘original sine wave #2’ in Beattie (2015), and is likely to be impractical or uninformative for real experiments.

**Figure 19.**
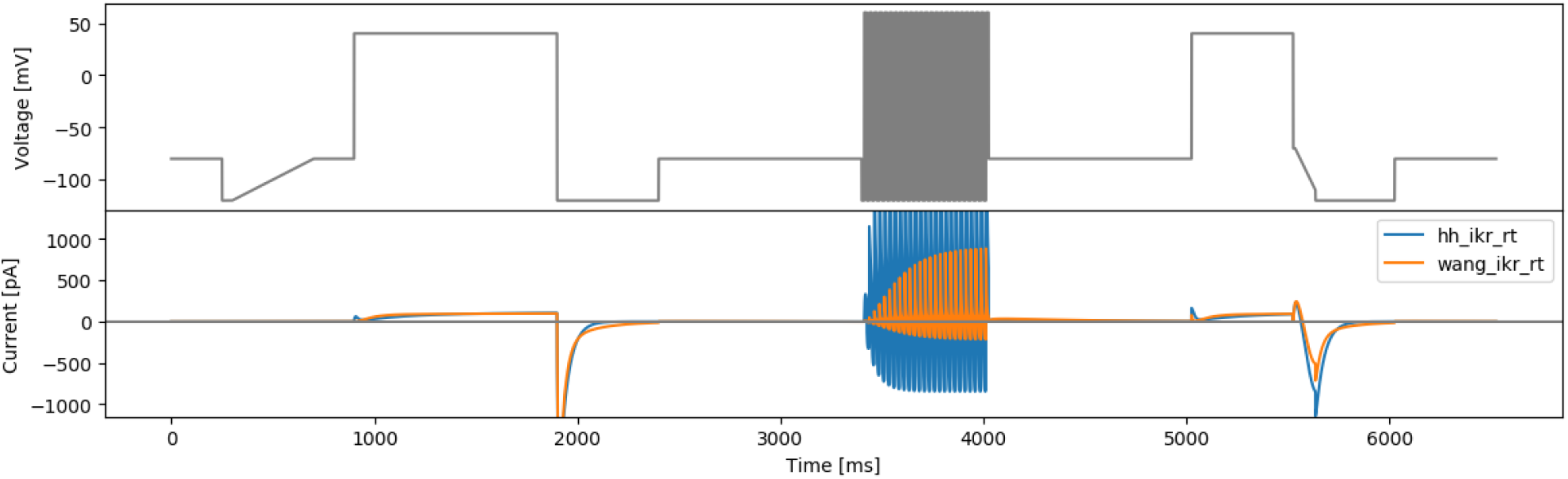
The square wave protocol for maximising two models’ difference (maxdiff) and simulated currents from both models.

## 7 Discussion

Developing ion channel models remains a challenging task predominantly due to all the various sources of uncertainty and variability (Mirams et al., 2016) — in terms of modelling approximations (Lei et al., 2020c; Lei and Mirams, 2021) as well as experimental noise and artefacts (Lei et al., 2020a). It is made more difficult due to the sparsity of available data for independent training and validation, with it still being common to calibrate models to all available data (Whittaker et al., 2020). The protocols presented here encompass many design criteria, including parameterisation, model selection and rigorous testing of the underlying assumptions in hERG models (Fink and Noble, 2009; Lei et al., 2019b; Mirams et al., 2024). As such, we expect that this collection of voltage clamp protocols will be extremely useful for development of mathematical models for the physiological gating of the hERG potassium channel, and in particular by providing ample validation data for assessing their prediction errors due to model discrepancy (Shuttleworth et al., 2024).

The same design criteria we have outlined here could easily be applied to other ion channels to create similar suites of protocols, using the provided open source codes.

## Data Availability

Open source code to reproduce the protocols in this report can be found on GitHub: https://github.com/CardiacModelling/protocol-design-hERG

## Acknowledgements

This work was supported by the Wellcome Trust (grant no. 212203/Z/18/Z); the Science and Technology Development Fund, Macao SAR (FDCT) [reference no. 0155/2023/RIA3 and 0048/2022/A]; the University of Macau [reference no. SRG2024-00014-FHS and FHS Startup Grant]; the Engineering & Physical Sciences Research Council (EPSRC) [grant no. EP/R014604/1]; and the Australian Research Council [grant no. DP190101758]. DGW & GRM acknowledge support from the Wellcome Trust via a Wellcome Trust Senior Research Fellowship to GRM. CLL acknowledges support from the FDCT and support from the University of Macau. We acknowledge Victor Chang Cardiac Research Institute Innovation Centre, funded by the NSW Government. The authors would like to thank the Isaac Newton Institute for Mathematical Sciences for support and hospitality during the programme The Fickle Heart when some work on this report was undertaken, and in particular discussions with Prof. Ian Vernon.

This research was funded in whole, or in part, by the Wellcome Trust [212203/Z/18/Z]. For the purpose of open access, the authors have applied a CC-BY public copyright licence to any Author Accepted Manuscript version arising from this submission.

## Notes

https://github.com/CardiacModelling/protocol-design-hERG

### Competing Interest Statement

The authors have declared no competing interest.

